# TCRanalyzer: A user-friendly tool for comprehensive analysis of T-cell diversity, dynamics and potential antigen targets

**DOI:** 10.1101/2025.05.09.652820

**Authors:** Nicole Seifert, Sarah Reinke, Nadine S. Kurz, Jakob Demmer, Kevin Kornrumpf, Tim Beißbarth, Wolfram Klapper, Michael Altenbuchinger

**Affiliations:** Department of Medical Bioinformatics, University Medical Center Göttingen, Germany; Department of Pathology, Hematopathology Section and Lymph Node Registry, University Hospital Schleswig-Holstein, Kiel, Germany; Campus Institute for Data Science (CIDAS), Section of Medical Data Science (MeDaS), University of Göttingen, Germany

## Abstract

T cells are critical for immune responses, recognizing antigens via their unique T-cell receptors (TCRs). Analyzing the diverse TCR repertoires, especially the hypervariable CDR3 region, is essential for understanding immune function in health and disease. Current TCR analysis tools often require specialized expertise, computational resources, or sacrifice biological information for efficiency. To address these limitations, we developed *TCRanalyzer*, a fast and comprehensive TCR analysis pipeline within a user-friendly graphical interface. *TCRanalyzer* covers all steps from data loading, aggregation and optional sequence clustering, to the analysis of TCR diversity metrics, clonal expansion and antigen specificity. Applied to datasets from patients with either benign or malignant tumors, *TCRanalyzer* identified changes in TCR clonality, clonal expansion and shifts in antigen specificity across different cohorts or following immunotherapy, thereby demonstrating its potential to dissect critical immunological processes. *TCRanalyzer* provides a robust and user-friendly tool for TCR sequence analysis, enhancing research in immunology and related fields.

**Availability:** *TCRanalyzer* is available at https://hub.docker.com/r/tcranalyzer/application.

## 1 Introduction

T cells play an important role in effective immune responses against various diseases by identifying and responding to antigens presented to their unique T-cell receptors (TCR). TCRs determine the antigen specificity of T cells through their hypervariable regions, such as the Complimentary Determining Region 3 (CDR3). The high variability of CDR3 contributes to the vast diversity and size of TCR repertoires, comprising up to billions of unique TCRs in individuals.

Advancements in high-throughput sequencing technologies offered cost-effective and scalable solutions for research purposes, enabling a sensitive detection of CDR3 sequences across diverse biological samples. In parallel with technological advancements, computational tools have played an essential role to gain insights from CDR3 sequencing data. Notable computational tools, such as *VDJtools* or *MiXCR*, offer a variety of functionalities that are tailored to different research questions and experimental designs, including the quantification of TCR diversity metrics, identification of clonally expanded T-cell populations, or inference of antigen-specific responses.^1,2^

Despite the availability of various tools, standardized pipelines for downstream analysis that are both user-friendly and comprehensive remain rare. Many existing tools require specialized expertise or high-performance computing resources, posing challenges for researchers with limited expertise and resources.^3,4^ Additionally, some tools sacrifice valuable biological information for computational efficiency, raise data security concerns by requiring the upload of sensitive data to external servers, or are non-operational due to a lack of maintenance.^5,6^

Addressing these challenges, we developed *TCRanalyzer*, a fast and sensitive TCR analysis pipeline that integrates famous tools like *immunarch* for data processing or *MMseqs2* for fast sequence clustering and alignment. *TCRanalyzer* provides a comprehensive analysis of TCR repertoires in a graphical user interface with only four concise modules. It is designed for ease of use and local deployment, making it accessible to a broad range of researchers and clinicians.

## 2 Methods

### 2.1 Overview

*TCRanalyzer* consists of four modules (see figure 1) designed to cover the entire workflow from raw data to in-depth analysis. Each module can be executed independently, providing flexibility to the user.

**Figure 1.**
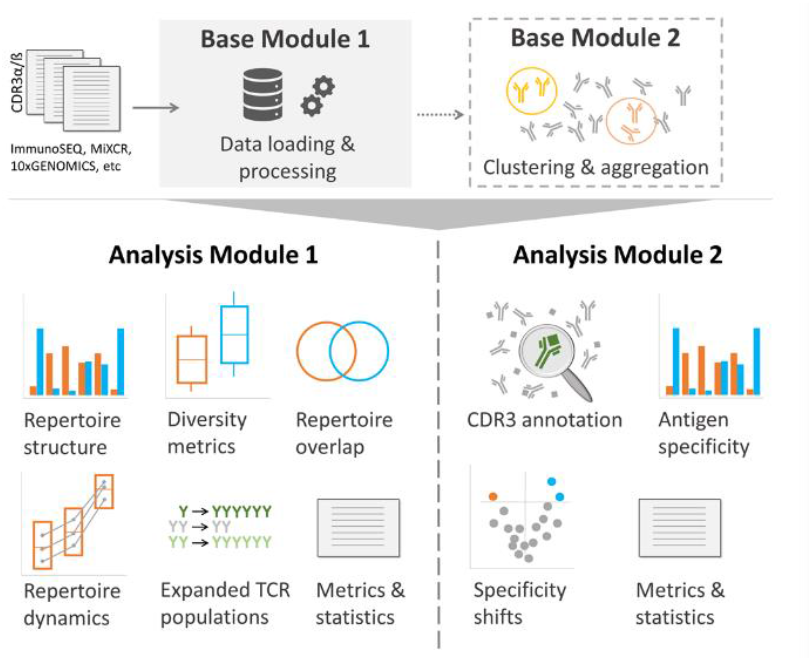
Overview of the functionalities of *TCRanalyzer’s* data preprocessing (Base Module 1, 2) and analysis modules (Analysis Module 1, 2)

### 2.2 Base Module 1: Data Loading and Initial Aggregation

Base Module 1 is essential for loading and transforming various sequencing data formats into a universal format and aggregating CDR3 sequences of the same amino acid clonotype. This module must be performed once for each new dataset before any other module can be started.

First, data are loaded and converted into a universal format using the R package *immunarch*.^7^ Currently supported data formats include all popular TCR analysis and post-analysis formats, such as those from *immunoSEQ, MiXCR, VDJtools, 10Xgenomics* and more (see https://immu-narch.com/ for more information). After conversion, datasets are split into subsets according to sample name specifics (see ReadMe, chapter VI). This step is necessary to allow for both individual and comparative analyses in subsequent Analysis Modules. Within each sample, all CDR3 sequences of a sample that are identical on amino acid level are then aggregated by summing up their counts. This is done to reduce dimensionality in the data, focusing on T-cell populations and their response to shared targets rather than on individual T-cell origins. The final datasets are stored in the provided data folder using the R package *qs* for quick saving and reading.^8^

### 2.2 Base Module 2: Clustering of CDR3 Sequences (Optional)

Base Module 2 addresses the challenge of high diversity of CDR3 sequences that can target the same antigen by clustering and aggregating CDR3 with similar amino acid sequences. This module is optional, but must be performed once for each new dataset before any Analysis Module can be started with clustered datasets.

This module loads all files generated in Base Module 1 and employs the easy-cluster algorithm of *MMseqs2* to perform cascaded clustering on all CDR3 sequences of each subset.^9^ Within each sample, CDR3 sequences of the same cluster are aggregated by summing up all respective counts and assigning the leading sequence of the cluster as the new CDR3 sequence. The clustered and aggregated datasets are stored separately in the provided data folder as done in Base Module 1.

### 2.4 Analysis Module 1: Clonality and Expansion Analysis

Analysis Module 1 focuses on the structure, diversity and dynamics of TCR repertoires and provides comparative analyses of different groups (e.g. different subsets). The input datasets for this module are selected by the user, choosing between the aggregated datasets from Base Module 1 or the further clustered datasets from Base Module 2.

In a first step, four metrics are calculated within each subset to describe the proliferative activity of T cells: Simpson’s Clonality (SC), Percentage of Singletons (PoS) and clonal expansion of Singletons (SE) and Non-Singletons (NSE). SC is a measure for clone homogeneity with higher values indicating a lower diversity within a given sample. SC is calculated for each sample as 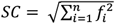, where *n* is the total number of different clones and *f*_*i*_ is the productive frequency of clone *i*. PoS represents the proportion of clones that are detected only once (Singletons) in a given sample. Clonal expansion is calculated per patient, if at least two samples of the same patient, collected at different timepoints, are available. Clonal expansion is calculated as previously described,^10^ but for each available time interval separately instead of restricting to the maximum expansion value. SE and NSE represent the percentage of expanded Singletons and Non-Singletons, respectively.

In a second step, this module allows for comparison of these metrics between two different subsets (e.g. different diseases or T-cell subtypes) or timepoints (e.g. before and after therapy) as initially selected by the user (see ReadMe, chapter IV.4). It performs statistical tests, automatically selecting the appropriate test based on whether the data is paired or not. Additionally, this module calculates the percentage of overlapping sequences between the two groups. It also uses the functions of the R package *immunarch* to visualize differences in (a) CDR3 lengths, (b) relative abundance of rare and hyperexpanded T-cell populations, (c) number of unique CDR3 sequences and (d) V gene usage between the two groups (see https://immunarch.com/ for more information). Calculation of (a)-(c) is based on aggregated or clustered CDR3 amino acid sequences, as chosen initially.

Analysis Module 1 can be repeated with different parameters for the comparative analyses in step 2. The computing time for additional comparative analyses will be reduced as calculation of the four metrics is done only once for a new input dataset and results are stored in the provided data folder as done in Base Module 1.

Outputs of Analysis Module 1 include (i) PDF files visualizing the metrics within a subset and the comparisons between groups, (ii) Excel files with calculated metrics per sample and test statistics, and (iii) RDA files of generated plots for further adjustment in R (see ReadMe, chapter VII for details on plot manipulation).

### 2.5 Analysis Module 2: Antigen Specificity

Analysis Module 2 focuses on CDR3-antigen annotation and a comparative analysis of antigen frequencies. The input datasets for this module are selected by the user, choosing between the aggregated datasets from Base Module 1 or the further clustered datasets from Base Module 2.

In a first step, all samples of all input datasets (subsets) are annotated and antigen frequencies are calculated. This module first combines all unique CDR3 sequences of all samples of a subset (input CDR3s). As initially selected by the user, input CDR3s are then aligned against either all CDR3α or all CDR3β sequences of the CDR3-antigen database *VDJdb* (db CDR3s) using the easy-search algorithm of *MMseqs2* (see ReadMe, chapter IV.6).^9,11^ For each input CDR3, all significant alignments with db CDR3s (E-value ≤ 0.05) are further filtered for the best unique alignment, starting with lowest E-value and continuing with highest bit-score and highest sequence identity, if several alignments show identical values. For each remaining alignment, the antigen annotation of the respective db CDR3 is extracted. If an input CDR3 has several antigen annotations that are not identical, the input CDR3 is either not annotated (Annotation: *exactly*) or the most frequent antigen is annotated (Annotation: *max*; see ReadMe, chapter IV.7 for more details). Input CDR3s with no significant alignment receive no annotation. Finally, this module calculates the antigen frequencies within a sample, with each frequency reflecting the proportion of the total annotated CDR3 sequences that are annotated with a specific antigen.

In a second step, this module allows for comparison of antigen frequencies between two different subsets (e.g., different diseases or T-cell sub-types) or timepoints (e.g., before and after therapy) as initially selected by the user (see ReadMe, chapter IV.4). It performs statistical tests, automatically selecting the appropriate test based on whether the data is paired or not, and calculates the mean-fold-change in antigen frequencies between both groups.

Analysis Module 2 can be repeated with different parameters for the comparative analyses in step 2 (see ReadMe, chapter IV). The computing time for additional comparative analyses will be reduced as the alignment, annotation and calculation of antigen frequencies in step 1 are done only once for a new input dataset and results are stored in the provided data folder as done in Base Module 1.

Outputs include (i) PDF files of volcano plots visualizing differences in antigen targets between groups, (ii) Excel files with antigen frequencies and test statistics, and (iii) RDA files of generated plots for further adjustment in R (see ReadMe, chapter VII for more details on plot manipulation).

## 3 Results

A case study was conducted using TCR repertoires of 8 patients with benign lymph nodes (BLN) and publicly available data of 14 patients with hepatocellular carcinoma (HCC) undergoing immunotherapy.^12^ For each patient, TCR repertoires were available at initial diagnosis (T0) and one follow-up timepoint (T1; BLN: without treatment, HCC: after immunotherapy). TCR repertoires were collected from the affected tissue of each patient and additionally for each HCC patient, also from the peripheral blood. Therefore, the input dataset for *TCRanalyzer* consisted of 72 samples, which were assigned to three subsets called BLN, HCCtissue and HCCblood. Data is provided as an example dataset for *TCRanalyzer* at https://gitlab.gwdg.de/MedBioinf/immunology.

*TCRanalyzer* efficiently processed and analyzed the data, providing insights into differences between subsets, and also into TCR repertoire changes in HCC following immunotherapy.

Analysis Module 1 revealed significant differences in Simpson’s Clonality, the percentage of singletons and clonal expansion of Non-Singletons between benign (BLN) and malignant tumors (HCCtissue). It was also able to uncover divergent T-cell activity in the tumor microenvironment and the peripheral blood of HCC patients (HCCtissue vs HCCblood). The results indicate an increased proliferative activity of T cells in malignant tumors, already at timepoint of diagnosis. They also show further clonal expansion of these T-cell populations following immunotherapy, indicating that immunotherapy reactivated pre-expanded clones rather than activating new ones.

Analysis Module 2 revealed divergent antigen specificities of T cells in the tumor microenvironment and the peripheral blood of HCC patients (HCCtissue vs HCCblood). Even though annotations of neoantigens in *VDJdb* are rare, *TCRanalyzer* was able to detect an increase in PABPC1 specificity, a gene known to be prognostic in HCC, in the peripheral blood of HCC patients following immunotherapy.^13,14^

## 4 Discussion

*TCRanalyzer* addresses the limitations of traditional TCR analysis methods, offering a comprehensive analysis of TCR repertoires through a user-friendly Dockerized Shiny App interface. The integration of *immunarch* and *MMseqs2* provides robust data processing and very fast clustering and alignment capabilities. *TCRanalyzer* is able to quantify TCR diversity metrics, track dynamic changes and identify clonally expanded T-cell populations. Integrating the CDR3-antigen database *VDJdb, TCRanalyzer* is also able to infer antigen-specific responses and investigate differences over time or across cohorts. Unfortunately, antigen annotation is currently limited by the numbers of annotated CDR3 sequences in *VDJdb*, especially with regard to neoantigens. The inclusion of more comprehensive CDR3-antigen databases could improve *TCRanalyzer*’s utility in the future.

## Supporting information

ReadMe v2

## Data Availability

TCR repertoires from 14 HCC patients undergoing immunotherapy are publicly available under Digital Object Identifier https://doi.org/10.21417/RP2024NM. TCR repertoires from 8 patients with BLN are available upon request.

## Authorship Contributions

NS, SR, MA and WK conceived and designed the overall analysis pipeline. NS, NK, JD and KK developed the Shiny App Interface. NS, SR, JD and WK generated and analyzed data. NS wrote the manuscript. All authors revised the manuscript and agreed with its final version.

