## Supplement for "TCRanalyzer: A user-friendly tool for comprehensive analysis of T-cell diversity, dynamics and potential antigen targets"

### Supplementary Figures

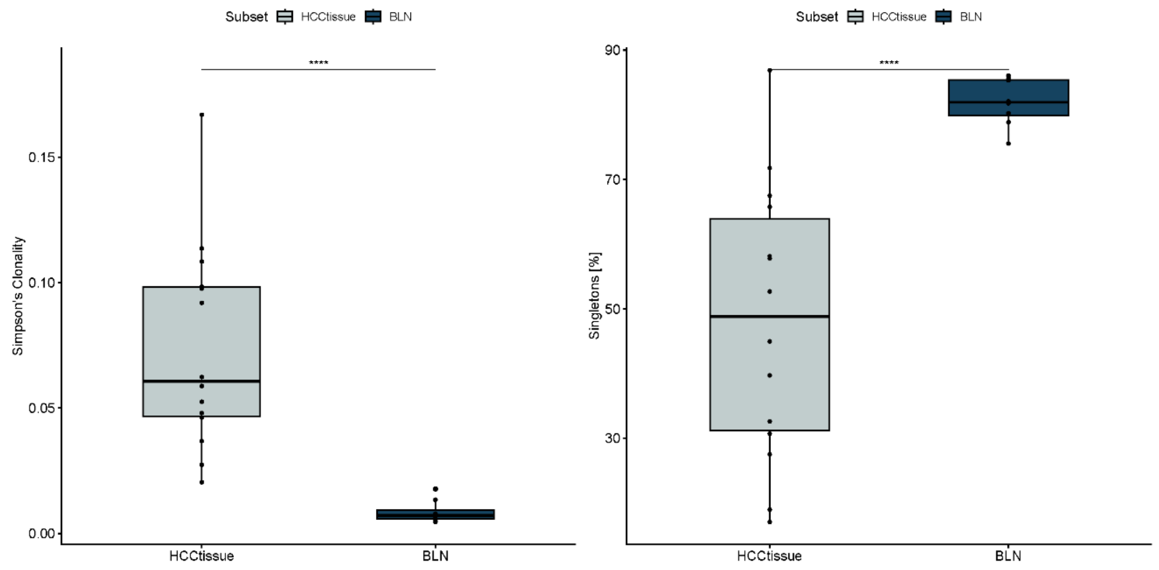


**Figure 1**: Simpson's Clonality (left) and Percentage of Singletons (right) in tissue biopsies of patients with hepatocellular carcinoma (HCCtissue) or with benign lymph nodes (BLN) at initial diagnosis (T0).

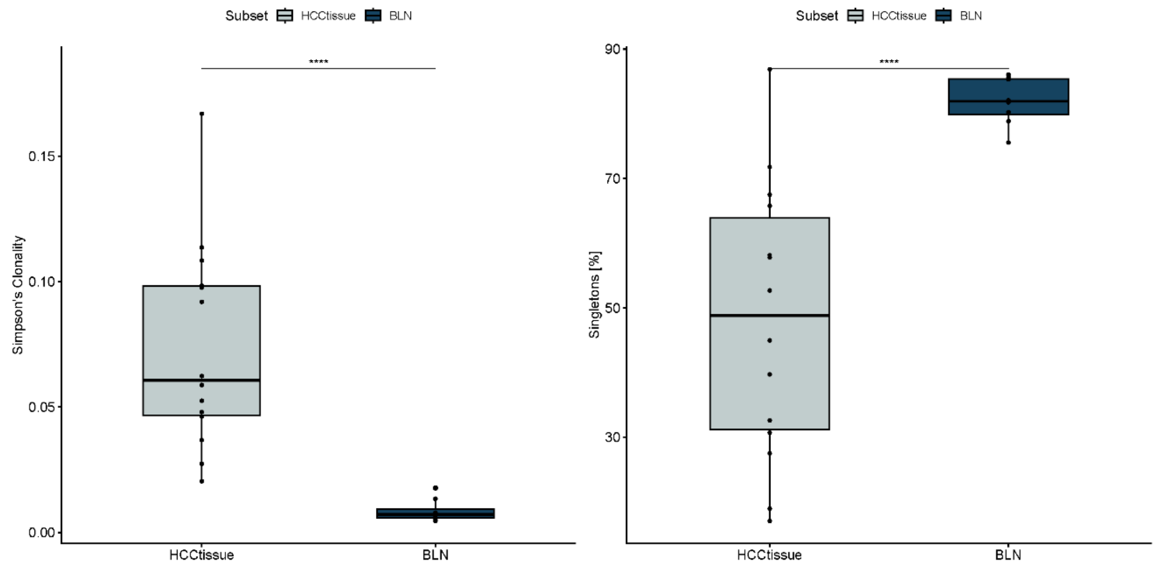


**Figure 2**: Simpson's Clonality (left) and Percentage of Singletons (right) in tissue biopsies of patients with hepatocellular carcinoma (HCCtissue) after immunotherapy or with benign lymph nodes (BLN) at a follow-up diagnosis (T1).

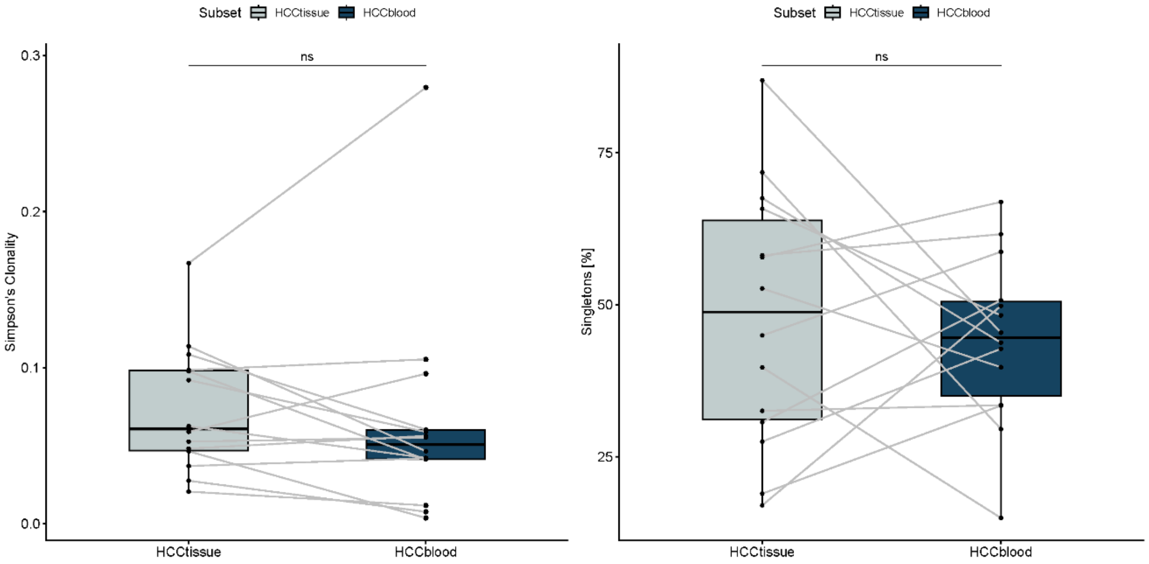


**Figure 3**: Simpson's Clonality (left) and Percentage of Singletons (right) in paired tissue biopsies (HCCtissue) and samples of peripheral blood mononuclear cells (HCCblood) of patients with hepatocellular carcinoma at initial diagnosis (T0).

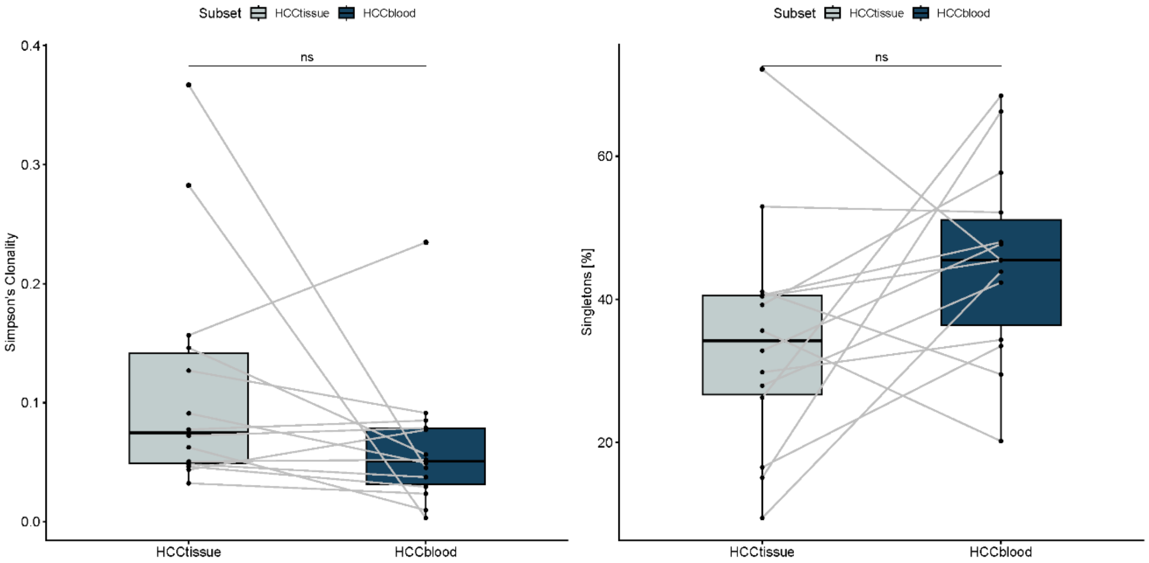


**Figure 4**: Simpson's Clonality (left) and Percentage of Singletons (right) in paired tissue biopsies (HCCtissue) and samples of peripheral blood mononuclear cells (HCCblood) of patients with hepatocellular carcinoma after immunotherapy (T1).

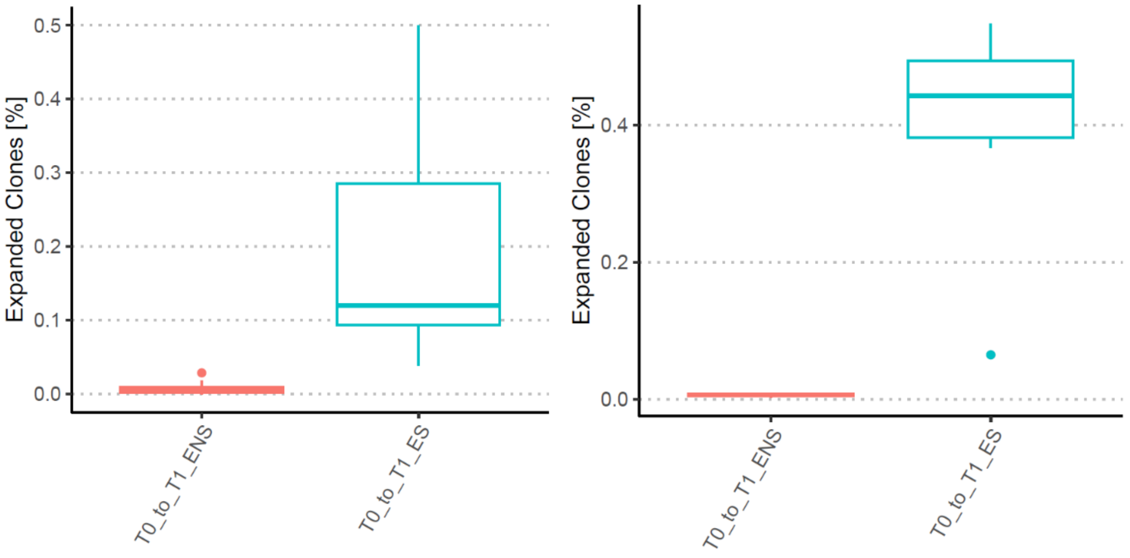


**Figure 5**: Expansion of Non-Singletons (ENS) and of Singletons (ES) from T0 (initial diagnosis) to T1 (after immunotherapy) in paired tissue biopsies (left) and samples of peripheral blood mononuclear cells (right) of patients with hepatocellular carcinoma.

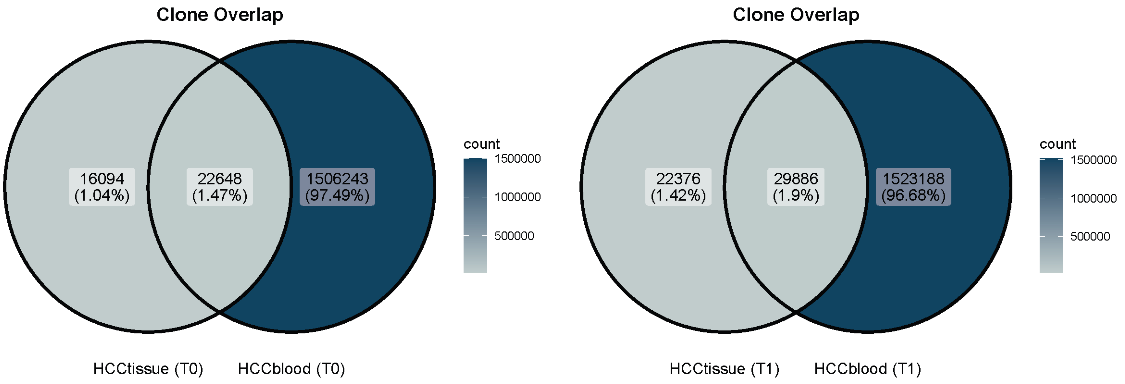


**Figure 6**: Overlap of unique CDR3 sequences in paired tissue biopsies (HCCtissue) and samples of peripheral blood mononuclear cells (HCCtissue) of patients with hepatocellular carcinoma at initial diagnosis (T0, left) and after immunotherapy (T1, right).

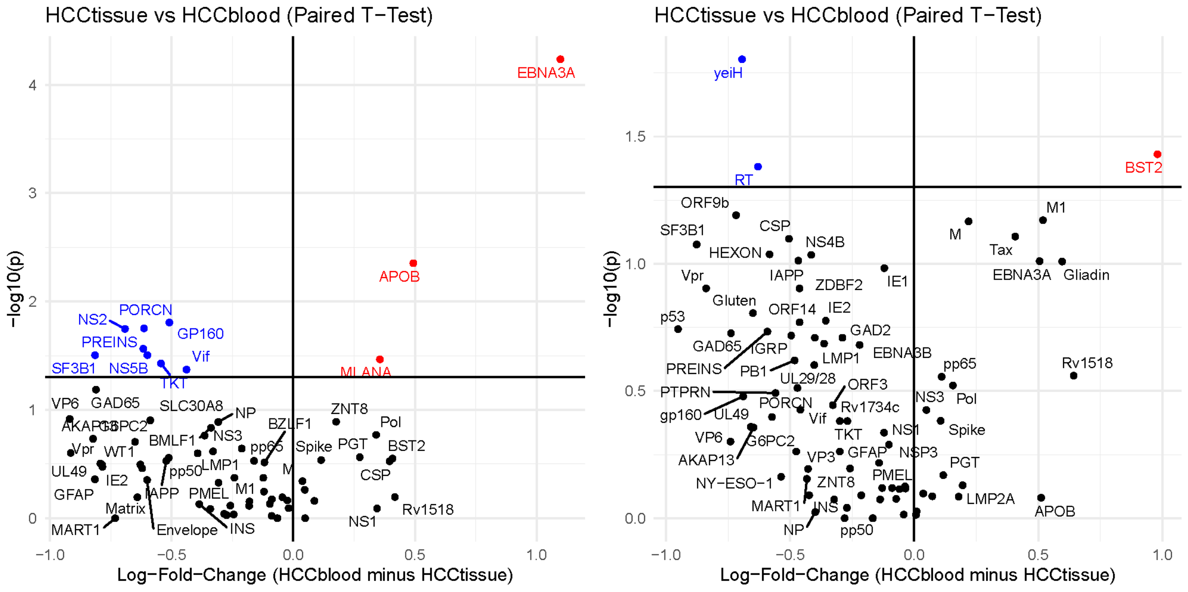


**Figure 7**: Antigen specificities of CDR3 sequences compared between paired tissue biopsies (HCCtissue) and samples of peripheral blood mononuclear cells (HCCtissue) from patients with hepatocellular carcinoma at initial diagnosis (T0, left) and after immunotherapy (T1, right).
