## Supplementary material for "TCRanalyzer: A user-friendly tool for comprehensive analysis of T-cell diversity, dynamics and potential antigen targets": ReadMe v2

T cells are critical for immune responses, recognizing antigens via their unique T-cell receptors (TCRs). Analyzing the diverse TCR repertoires, especially the hypervariable CDR3 region, is essential for understanding immune function in health and disease. Current TCR analysis tools often require specialized expertise, computational resources, or sacrifice biological information for efficiency. To address these limitations, we developed `TCRanalyzer`, a fast and comprehensive TCR analysis pipeline within a user-friendly graphical interface. `TCRanalyzer` covers all steps from data loading, aggregation and optional sequence clustering, to the analysis of TCR diversity metrics, clonal expansion and antigen specificity. Applied to datasets from patients with either benign or malignant tumors, `TCRanalyzer` identified changes in TCR clonality, clonal expansion and shifts in antigen specificity across different cohorts or following immunotherapy, thereby demonstrating its potential to dissect critical immunological processes. `TCRanalyzer` provides a robust and user-friendly tool for TCR sequence analysis, enhancing research in immunology and related fields.

---

#### Table of Contents


- I. Installation
- II. Overview
- III. Quick start
- IV. Parameters
- V. How to use your own data
- VI. Plot manipulation
- VII. Troubleshooting
- VIII. Contact
- IX. License

### I. Installation

TCRanalyzer is available out-of-the-box as a Docker container (https://hub.docker.com/r/tcranalyzer/application), which may be started within a graphical environment (Docker Desktop) or from the command line.

#### Option 1 (recommended): Use Docker Desktop

1. Start Docker Desktop (Download available at: https://www.docker.com/products/docker-desktop/)
2. Search for 'tcranalyzer/application' and click 'Pull'

Wait until the download is finished and that's it. You can now follow the Quick start guide below to start your first analysis in `TCRanalyzer`.

#### Option 2: Command line installation (Powershell, Bash)

To use TCRanalyer from the command line, make sure that Docker is installed (https://docs.docker.com/engine/install/).

Pull the TCRanalyzer Docker container from Docker Hub:

```
docker pull tcranalyzer/application:latest
```

#### II. Overview

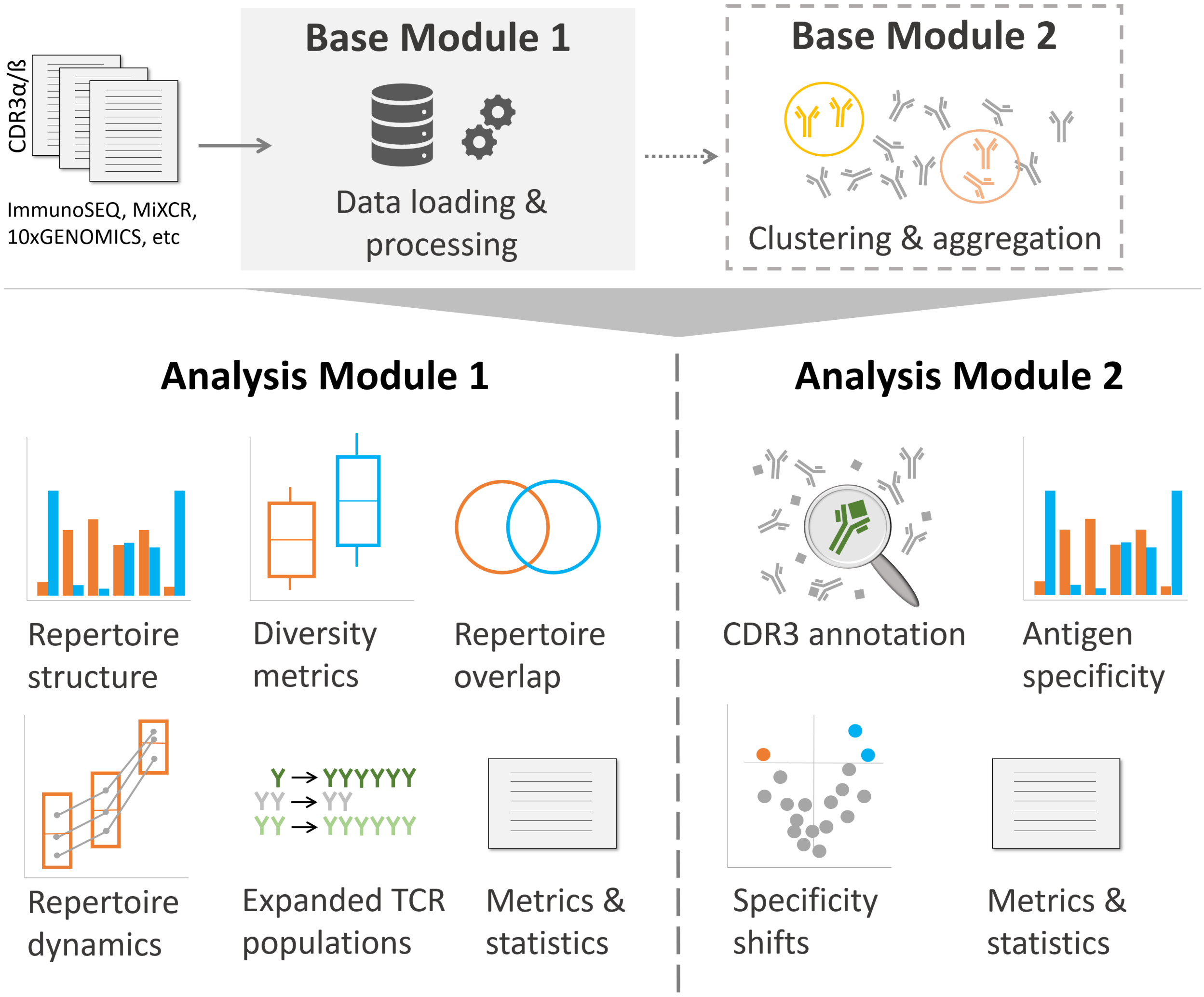
CDR3α/ß
ImmunoSEQ, MiXCR, 10xGENOMICS, etc
Base Module 1
Data loading & processing
Base Module 2
Clustering & aggregation
Analysis Module 1
Repertoire structure
Diversity metrics
Repertoire overlap
Repertoire dynamics
Expanded TCR populations
Metrics & statistics
Analysis Module 2
CDR3 annotation
Antigen specificity
Specificity shifts
Metrics & statistics

### III. Quick start

**Important:** Before starting TCRanalyzer, always ensure your sample names follow the proper format (“**PatID_SubsetX_Tx**”) and that your sequencing data is stored in a folder structured as `MyFolder/studyname/`. Please refer to our How to use your own data guide for more information.

#### 1) Start the Docker container

We recommend using Docker Desktop on a Windows machine. However, you can also start Docker container using the command line (Powershell, Bash). More details are provided at the end of this Quick Start guide (section 5) Optional).

**Use Docker Desktop**

**1. Run "tcranalyzer/application" in Docker Desktop**

- Search for "tcranalyzer/application" (in "Images")
- Click "Run" (in "Actions")

**2. Add "optional settings"**

- Host port = 3838,
- Host path = select the folder you created (e.g. named "MyFolder", which contains one or more folders with the sequencing data you want to analyze, e.g. named "studynames")
- Container path= /app/Data

#### 2) Open the app

Open `http://localhost:3838` in your webbrowser

Note: By accessing "localhost" your data remains on your local computer and analyses are performed on your own system. It will not be forwarded to the internet.

#### 3) Start your analyses

1. Choose a module
2. Select or fill in parameters
3. Click Run

**That's it, TCRanalyzer does the rest. You can find a folder containing all generated results in the folder you provided in step 1.**

Please read our Paper for a detailed description of the functionalities of each module.

Note: Run Base Module 1 for every new dataset (and optional: Base Module 2 afterwards). Analysis Modules 1 & 2 build upon the outputs of the Base Module(s).

#### 4) Close TCRanalyzer

You can stop the app in Docker Desktop (Container -> Actions -> Stop)

#### 5) Optional:

**Start the Docker container using the command line (Powershell, Bash)**

Start the container by mounting your desired directory into the directory `/app/Data` within the TCRanalyzer Docker container. This mounted volume becomes your working environment. Within it, a **Data** folder and a **Results** folder will be created automatically when you run the modules.

Below is an example command with explanations for each flag:

```
docker run -it --rm -p 3838:3838 -v /MyPath/MyFolder:/app/Data -u `id -u $USER` --name tcranalyzer tcranalyzer/application:latest
```

**Explanation of flags:**

- **`-it`**
  Runs the container in interactive mode with a TTY attached. It is useful for debugging with real-time output.

- **`--rm`**
  Automatically removes the container when it exits.

- **`-p 3838:3838`**
  Maps port **3838** of the container to port **3838** on the host. This allows you to access the Shiny application through your browser using `http://localhost:3838`.

- **`-v /MyPath/MyFolder:/app/Data`**
  Mounts the host directory `/MyPath/MyFolder` into the container at `/app/Data`.
  **Note:** This mounted volume is your working environment. You need to place your sequencing data folder inside this mounted volume so that the modules can find and process the data correctly. When the container starts, it will create a **Data** folder and a **Results** folder within `/app/Data`.

- **`-u \`id -u $USER`** `Runs the container as the current host user by fetching the user ID using` id -u $USER`. This helps avoid file permission issues when writing to the mounted volume.

- **`--name tcranalyzer`**
  Assigns the name `tcranalyzer` to the container.

- **`tcranalyzer/application:latest`**
  Specifies the Docker image to use (with the `latest` tag). Make sure this image exists on your system or in your repository.

With this setup, after you start the container, your working environment is set up in `/MyPath/MyFolder` (mounted as `/app/Data` inside the container). The modules will create and populate the **Data** and **Results** folders within that directory based on your analysis workflows.

###### Backup specific files (generating ZIP file)

In the Backup tab you can select specific folders and files from your working environment to create a ZIP archive. This ZIP file is generated and downloaded locally to your PC. Nothing is stored externally - sensitive data remains on your system.

- **File Selection:**
  Use the interactive folder tree to choose the folders and files you want to include in the backup.
- **Filename:**
  Enter your desired filename in the provided text field. The application will create a ZIP file with that name.
- **ZIP Creation:**
  Click the "Zip and Save Selected Data for a Backup" button. The selected data is compressed into a ZIP file, which is then downloaded directly to your PC.

By performing the backup locally, your data stays secure on your machine and is never transmitted over the web.

###### Load zipped Data into your Working Environment

The **Load Study Files** tab in TCRanalyzer lets you import your study data directly from a ZIP file into your working environment - no manual file copying or external uploads required. Here's how it works:

- **Study Name Assignment:**
  Enter a study name in the provided text field. This name becomes the folder name in your working environment. All data extracted from the ZIP file will be organized under this folder.
- **ZIP File Selection:**
  Use the file input to browse and select your ZIP file. The selected ZIP file contains your study's sequencing data. Importantly, the file is not uploaded to any remote server - it remains local and is only used to extract its contents into the container’s mounted volume.
- **Data Extraction:**
  When you click the "Extract Study Data" button, the application will unzip the file directly into your working environment (the mounted volume, typically found at `/app/Data`). During extraction, two folders are created automatically: one for raw data (Data) and one for analysis results (Results).
- **Local Working Environment:**
  The mounted volume serves as your working environment, where all study data is stored and processed. This approach ensures that your data stays on your local system (or in your designated workspace) and is immediately available for analysis by the container’s modules.

This feature streamlines the process of importing study data. Instead of manually copying and extracting files via your file browser, you simply select the ZIP file in the UI, and the data is automatically extracted into your working directory. This makes it easier and more efficient to update or import new data without leaving the TCRanalyzer interface.

### IV. Parameters

#### 1) Study name

= folder name containing your sequencing data

- This parameter is needed in every module and tells TCRanalyzer which dataset you want to use
- Example: HCC

#### 2) Count cutoff

= All samples with a total CDR3 count < cutoff will be removed from the dataset.

- Default: 100 (can be changed to any other number)

#### 3) Clustered TCR

= Do you want to analyze unclustered ("No") or clustered CDR3 ("Yes")?

- "No": aggregated datasets from Base Module 1 will be analyzed
- "Yes": aggregated & clustered datasets from Base Module 2 will be analyzed
- Note: If you select "yes" and get an error, please check if you ran Base Module 2 on the dataset once.

#### 4) Comparison

Do you want to compare TCR repertoires between two subsets at a defined timepoint ("Subset") or between two timepoints in a subset ("Timepoint")?

- "Subset": TCRanalyzer compares TCR repertoires of two different subsets at a defined timepoint
  - **Subset 1**: name of the first subset (e.g. "Healthy")
  - **Subset 2**: name of the second subset (e.g. "HCC")
  - **Timepoint**: timepoint of the samples of the chosen subsets (e.g. "T1")
- "Timepoint" TCRanalyzer compares TCR repertoires of the same subset, but between two different timepoints
  - **Timepoint 1**: first timepoint of the chosen subset (e.g. "T0")
  - **Timepoint 2**: second timepoint of the chosen subset (e.g. "T1")
  - **Subset**: name of the subset (e.g. "HCC")

##### 5) Compare dynamics?

Set **Compare Dynamics** to **Yes** if you want to analyze dynamic changes in TCR repertoires. Comparisons will be between specific time intervals. This option works differently depending on your chosen comparison mode:

**When Comparison = Subset**

You compare TCR repertoires of specific time intervals between two different subsets.

- **Time Interval 1:**
  The time interval for Subset 1 (e.g., "T0" and "T1" for "HCC").
- **Time Interval 2:**
  The time interval for Subset 2 (e.g., "T1" and "T2" for "Healthy").

**When Comparison = Timepoint**

You compare TCR repertoires of two specific time intervals within one subset (e.g. "HCC").

- **Time Interval 1:**
  The first time interval for the chosen subset (e.g., "T0" and "T1" for "HCC").
- **Time Interval 2:**
  The second time interval for the chosen subset (e.g., "T1" and "T2" for "HCC").

**Important Notes:**

- The available timepoints are parsed from the file names of your sequencing data that is available at runtime.
- If the timepoints of your desired comparison are not available please review the naming your sequencing data.
- If you have recently added or removed sequencing data please re-run atleast Base Module 1.
- Comparisons of empty time intervals are not supported (e.g., T0-T0 or T1-T1).

##### 6) TCR chain

Did you sequence CDR3 from TCR alpha ("TRA") or beta chain ("TRB")? Note: TCRanalyzer will select the compatible subset of CDR3 sequences in the CDR3-antigen database.

#### 7) Annotation

Decides on the type of antigen annotation that should be analyzed (recommended: gene.exactly). This parameter is needed for the comparison of antigen specificities between the two groups of TCR repertoires (as selected previously)

- "gene.exactly" (*recommended*): CDR3s are annotated with target genes, if the filtered alignments with VDJdb CDR3s have one or several *identical* target *gene* annotations
- "species.exactly": CDR3s are annotated with target species, if the filtered alignments with VDJdb CDR3s have one or several *identical* target *species* annotations
- "gene.max": CDR3s are annotated with the *most frequent* target *gene* in the filtered alignments with VDJdb CDR3s
- "species.max": CDR3s are annotated with the *most frequent* target *species* in the filtered alignments with VDJdb CDR3s

#### 8) Volcano Plot P-value Type

This parameter determines which type of p-value is used in the volcano plot. You can choose between:

- **p (corrected)**: Applies multiple testing correction using the Benjamini-Hochberg method to control the false discovery rate (FDR). This is **strongly recommended**, especially when working with large datasets where multiple comparisons increase the risk of false positives.
- **p**: Raw, unadjusted p-values. Use with caution, as these do not account for multiple testing and may lead to misleading results.

### V. How to use your own data

**Important**: If you change your dataset, you must either assign a new folder name for the updated dataset or delete the associated files in the Data and Results Directories, as these processes build upon one another's results. Please rerun at least BM1.

#### 1) Check sample names

- The names of your samples have to follow the format: "**PatID_SubsetX_Tx**"
- Important: Use only combinations of letter(s) and/or number(s) for **PatID** and **SubsetX** (special characters, symbols or spaces will result in errors)
- **PatID**: unique identifier for each patient, e.g. "Pat123"
- **SubsetX**: unique identifier for each subset, e.g. "Blood"
- **Tx**: unique identifier for timepoints, e.g. "T0" (use "T" followed by a number)
- Sample names could be: "Pat123_Blood_T0", "Pat123_Blood_T2", "Pat123_Tumor_T0", "Pat123_CD8_T3"

#### 2) Save your sequencing data in MyFolder/Data/studyname

- **MyFolder**: The folder where your data and all results of your analyses will be stored (We recommend creating a new folder for TCRanalyzer and save your recent and all future data for analysis there)
- **studyname**: To add new sequencing data for analysis, create a new subfolder in "MyFolder", e.g. "MyFolder/studyname" (Note: we recommend choosing a short, unique foldername without using any special characters or spaces. All generated results will carry this name.)
- Save only your sequencing data in "MyFolder/studyname", nothing else.
- Note: You can have several **studyname**-folders with different names. You will choose in each module which dataset yout want to use/analyze by providing the folder name (with parameter: Study name).

#### 3) Proceed with our Quick start guide

### VI. Plot manipulation

If you are familiar with R, you can try out the fixVis function of the R package 'immunarch' (https://immunarch.com/) to customize your plots. For this purpose, TCRanalyzer saves all generated plots as .rda files in: MyFolder/studyname/Results/AM1/fixVis (or /AM2/fixVis).

Dependencies:

- R & RStudio (How to install: https://rstudio-education.github.io/hopr/starting.html)
- Package immunarch (How to install: https://immunarch.com/)

#### 1) Open RStudio & load immunarch

```
library(immunarch)
```

#### 2) Load your plot

```
load("MyFolder/Results/studyname/AM1/fixVis/plotname.rda")
```

Or you can simply double-click on plotname.rda in the fixVis folder.

#### 3) Use the fixVis function of immunarch

```
fixVis(plot.tmp)
```

A graphical interface will open where you can adjust your plot manually.

Note: plot.tmp is the name of the plot in your global environment in R. You can exchange "plot.tmp" with another plot of your R environment.

You can visit https://immunarch.com/ for more information on plot manipulation with fixVis.

### VII. Troubleshooting

#### Sudden Stop of the Processes

**1. Unexpected Stopping in Virtual Machine (e.g., WSL):**

**Check if RAM is sufficient**: You can monitor the system's memory usage by running `htop` (if not installed, you can install it using `sudo apt install htop` in your virtual machine or Docker container). In `htop`, observe the memory and swap usage. If the system is running low on memory, it might cause processes to stop unexpectedly. If needed allocate more memory to your Virtual Machine.

**2. Unexpected Stopping in Docker Desktop**

**Allocate more RAM to the Docker container**: If you suspect insufficient memory is causing the issue in a Docker container, you can increase the allocated RAM by modifying the Docker settings:

- Open Docker Desktop.
- Go to **Settings** -> **Resources**.
- Adjust the **Memory** slider to allocate more RAM to the Docker container.

After adjusting the settings, restart the container to apply the changes.

#### Other Errors

In case of errors, please check the log files generated in the `Log` directory created in your Data directory.
TCRanalyzer generates a log file for each module, named `info_bm1.log`, `info_bm2.log`, `info_am1.log`, and `info_am2.log`. If you need help regarding your error, please contact us with the respective log file attached.

### VIII. Contact

You can create a ticket with your question or error on Gitlab Issues or contact us directly via.

### IX. License

The package is freely distributed under the CC-BY 4.0 International license.

For commercial or server use, please contact us via.
